# Illuminating endosomal escape of polymorphic lipid nanoparticles that boost mRNA delivery

**DOI:** 10.1101/2020.12.02.407601

**Authors:** Marco Herrera, Jeonghwan Kim, Yulia Eygeris, Antony Jozic, Gaurav Sahay

**Affiliations:** Department of Pharmaceutical Sciences, College of Pharmacy, Robertson Life Sciences Building, Oregon State University, Portland, Oregon, 97201, USA; Department of Biomedical Engineering, Robertson Life Sciences Building, Oregon Health & Science University, Portland, Oregon, 97201, USA; Department of Ophthalmology, Casey Eye Institute, Oregon Health & Science University, Portland, Oregon, 97239, USA

**Keywords:** Endosomal escape, Trafficking, mRNA delivery, Galectin-8, β-sitosterol

## Abstract

Lipid-based nanoparticles (LNPs) for the delivery of mRNA have jumped to the forefront of non-viral gene delivery. Despite this exciting development, poor endosomal escape after LNP cell entry remains an unsolved, rate-limiting bottleneck. Here we report the use of galectin 8-GFP (Gal8-GFP) cell reporter system to visualize the endosomal escaping capabilities of LNPs encapsulating mRNA. LNPs substituted with phytosterols in place of cholesterol exhibited various levels of Gal8 recruitment in the Gal8-GFP reporter system. In live-cell imaging, the results of the LNPs containing β-sitosterol (LNP-Sito) showed a 10-fold increase in detectable endosomal perturbation events when compared to the standard cholesterol LNPs (LNP-Chol), suggesting the superior capability of LNP-Sito to escape from endosomal entrapment. Trafficking studies of these LNPs showed strong localization with late endosomes. This highly sensitive and robust Gal8-GFP reporter system can be a valuable tool to elucidate intricacies of LNP trafficking and ephemeral endosomal escape events, enabling the advancement in gene delivery.

## INTRODUCTION

The fields of nanotechnology and gene delivery continue to co-evolve for the treatment of human diseases.^1–3^ Within this nanoverse, extensive efforts towards RNA based therapeutics have made this nanomedicine the leading genetic tool for gene replacement, overexpression, repair and/or knockdown.^2,4^ One of the leading non-viral vectors for successful delivery of such nucleic acids is lipid-based nanoparticles (LNPs) (**Fig. 1A**).^5,6^ The LNP landscape continues to be developed clinically as it shows its advantageous properties for the treatment of multiple diseases including gene knockdown therapy for inherited hepatic disorders,^6,7^ cancer,^8^ and more recently, rapid vaccine development for the ongoing SARS-CoV-2 pandemic.^9,10^

**Fig. 1.**
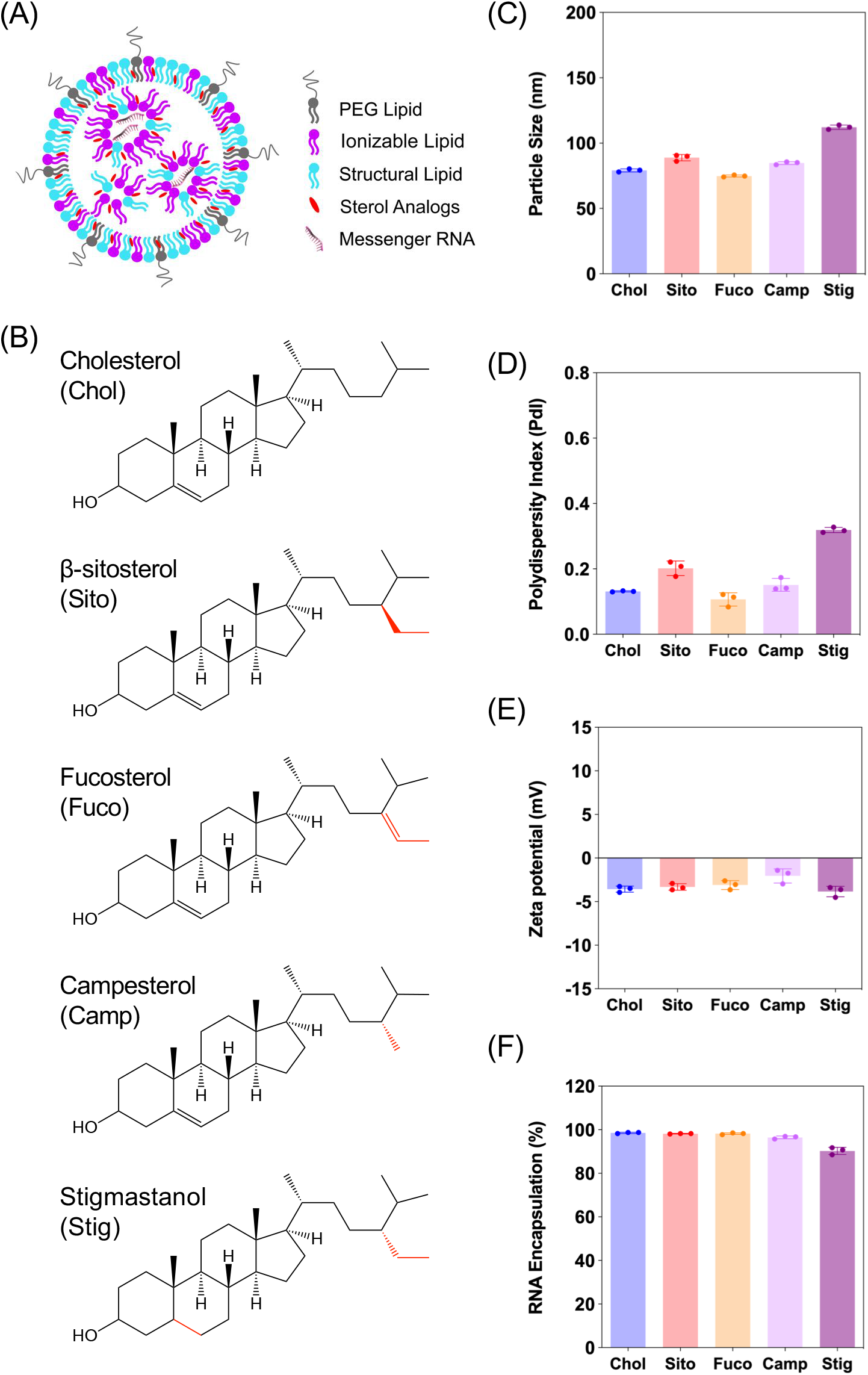
**(A)** Schematic illustration of a lipid-based nanoparticle (LNP) consisting of four lipids and messenger RNA encapsulated. **(B)** Chemical structures of the sterol analogs used in the present study: (top to bottom) Cholesterol (Chol), β-Sitosterol (Sito), Fucosterol (Fuco), Campesterol (Camp), and Stigmastanol (Stig). Structural differences are highlighted in red. **(C-E)** Physicochemical properties of LNPs containing various sterol analogs: **(C)** Particle size, **(D)** Polydispersity index (PdI) and **(E)** Zeta-potential of LNPs. **(F)** mRNA encapsulation of LNPs containing various sterol analogs.

The established mechanism of LNP-assisted intracellular gene delivery is via the evolutionarily conserved endosomal trafficking pathway.^11^ Although this cellular network has been extensively studied, there still are a lot of mechanistic minutiae that need to be uncovered to better understand and improve the targeting of this pathway for intracellular delivery. The process starts with interactions of LNPs with the plasma membrane they come in contact with, resulting in endocytosis. Once they are engulfed into the cell by endocytosis, these LNP-containing vesicles get taken along a continually evolving voyage that routes them to the early endosomes that act as the sorting hub. If not routed to the extracellular space via exocytosis or other organelles, the LNPs are transported to the multi-vesicular, late endolysosomal compartments. Throughout this complicated cellular odyssey, endosomally trapped LNPs are being subjected to a gradual drop in pH. Near the end of the late endosomal maturation stage, the LNPs undergo two major fates: recycling clearance by exocytosis from the cell^12^ or enzymatic degradation in lysosomes.

To exert therapeutic effects, exogenously delivered genes must be retrieved from the endolysosomal pathways to the cytosol;^11,13^ however, a minute fraction of particles (< 2%) escape into the cytoplasm following cell uptake.^14^ This underscores the importance of improving LNP endosomal escape to enhance potency. Despite great interest and efforts to breach the barriers faced, the exact mechanism of the endosomal escape of nanocarriers is limitedly understood. As more research focuses on understanding endosomal escape, the spotlight keeps expanding on the signaling players involved not only in the luminal endosomal system, but also in lysosomes and their closely-acting autophagosomes^15,16^. Nonetheless, investigating the endosomal escape of nanocarriers is extremely difficult because it is an ephemeral, low frequency event eliciting rapid disappearance; for example, escape of siRNA delivered by LNPs occurs within 5∼15 min of endocytosis.^17^ Current indirect visualization methods fail to give a clear picture of what is happening to the endosomes during these elusive endosomal disruption events.^14,18^ Therefore, improved visualization methods to illuminate the endosomal disruptions events following cell uptake are highly desired.

Galectins belong to a family of evolutionarily conserved β-galactoside-binding lectins found primarily diffused in the cytosolic compartment of cells^19,20^ and their biological functions include diverse pathways relating to cell homeostasis, cellular turnover, and immunity.^19–24^ One of the key biological functions is to sense damaged endosomes and accumulate specifically to the glycosylated inner leaflet of endosomal membranes.^25^ Out of several members of galectins, galectin 8 (Gal8) has emerged as a key sensor of damaged endosomes that also connects the closely ensuing autophagy.^23^ Recently, it has been exploited as a molecular indicator of the endosomal escape to crack the endosomal entrapment puzzle.^17^ Mechanistically, rapid recruitment of Gal8 to the damaged endosomes can be used to picture the intricacies responsible for the ephemeral endosomal escape events.^26,27^ Here, we employed a recently reported screening method based on a Gal8-GFP reporter fusion (Gal8-GFP) for the direct visualization of endosomal disruption events following cellular uptake of LNP-encapsulated mRNA into cells.

Utilizing the Gal8-GFP based direct fluorescence-guided assay, we report differences in endosomal escape capabilities elicited by varying sterol composition of LNPs used for the delivery of mRNA. We further elucidate the morphology of these LNPs, and correlate our findings to the bioactivity of the released cargo using live-cell imaging. Finally, we report colocalization patterns with late endosomal vesicles at the time of endomembrane disruption and attempt to connect the dots in this intricate machinery in hopes of advancing the endosomal escape elucidation for the delivery of gene therapies.

## EXPERIMENTAL

### Materials

Firefly luciferase (*Fluc*) mRNA fully substituted with 5-methoxyuridine was purchased from Trilink Biotechnologies (L-7202). (6Z,9Z,28Z,31Z)-heptatriacont-6,9,28,31-tetraene-19-yl 4-(dimethylamino)butanoate (DLin-MC3-DMA) was custom synthesized by Biofine International, Inc (BC, Canada). 1,2-distearoyl-sn-glycero-3-phosphocholine (DSPC) was purchased from Avanti Polar Lipids, Inc (Alabaster, AL). 1,2-Dimyristoyl-rac-glycero-3-methoxypolyethylene glycol-2000 (DMG-PEG_2K_), Cholesterol, β-Sitosterol, and Fucosterol were obtained from Sigma Aldrich (St Louis, MO). Campesterol and Stigmastanol were purchased from Cayman Chemicals (Ann Arbor, Michigan), and Matreya LLC (State College, PA), respectively. PB-GFP-Gal8 was a kind gift from Dr. Jordan Green (Addgene plasmid # 127191).

### LNP formulation and characterization

Lipid nanoparticles (LNPs) were prepared by microfluidic mixing using a previously described method.^28^ In short, lipid-ethanol solutions containing DLin-MC3-DMA, a sterol, DSPC, and DMG-PEG_2K_, at molar ratios of 50:38.5:10:1.5, were combined with 50 mM citrate buffer containing mRNA using a microfluidic mixer at a 1:3 ratio. LNPs were dialyzed twice using 3L of phosphate buffered saline (PBS, pH 7.4) and concentrated with 10 kDa Amicon® Ultra centrifuge filters (Millipore, Burlington, MA). Size distribution and zeta potential of LNPs were determined via dynamic light scattering using a Zetasizer Nano ZSP (Malvern Instruments, UK). mRNA encapsulation efficiency was determined using Quant-iT RiboGreen RNA reagent (Invitrogen, Carlsbad, CA).

### Development of the Gal8-GFP reporter cell line and culture

All Human Embryonic Kidney 293T/17 cells (ATCC CRL-11268, Manassas, VA) were cultured in Dulbecco’s Modified Eagle’s Medium supplemented with 10% heat-inactivated fetal bovine serum (Hyclone Laboratories Inc., Logan, UT) and 1X Penicillin/Streptomycin (Thermo Fisher, Federal Way, WA). All cultures were grown in 37 °C incubators supplemented with 5% CO2 and were cultured according to suppliers’ instructions. For stable transfections, Addgene plasmid #127191 coding for Piggybac Gal8-GFP as a 3’ terminus fusion protein was co transfected with Super Piggybac Transposase expression vector from System Biosciences (Palo Alto, CA) into 293T/17 cells. GFP positive cells were obtained following fluorescence activated cell sorting using BD FACSAria^™^ Fusion equipped with a 488nm laser and BD FACS Diva v8.0.1 software. Cells were sorted two separate times (Day 1 post transfection and day 8 post transfection), selecting for bright green fluorescence indicative of stable genomic integrations at AATT sites. All transfections and cell uptake studies were performed in passage 10-25 of the reporter cells after successful stable transfection were established.

### *In vitro* Fluc mRNA transfection assay

293T/17 cells were plated at 4,000 cells/well in white-wall, clear-bottom, 96 well plates. After overnight incubation, cells were treated with LNPs encapsulating *Fluc* mRNA at various doses. mRNA transfection results were collected at 3 hours or 24 hours after treatment using ONE-Glo^™^ + Tox luciferase reporter and cell viability kit (Promega, Madison, WI) with a multimode microplate reader. Luminescent signals (luciferase expression, RLU) were divided by fluorescent signals (cell viability, RFU) for normalization of the data according to the previous study.^28^

### Visualization of Gal8 recruitment in 293T/17 Gal8-GFP cells with treatment of various LNPs

Ibidi 8-well chamber slides (Fitchburg, WI) were coated with poly-D lysine (Thermo Fisher, Waltham, MA) and rinsed with PBS before seeding the 293T/17 Gal8-GFP reporter cells at 60,000 per well in complete media and allowed to adhere and incubate overnight. The next day, different LNPs were added dropwise onto the media at the concentration of 50 ng, 100 ng and 200 ng mRNA per well and incubated for 3 hr and 24 hr at which point the media was aspirated, cells were washed with PBS and fixed with 4% paraformaldehyde in PBS for 10 min at room temperature. After fixing, cells were washed with PBS three times and DAPI (ThermoFisher, Walthman, WA) was added at 1:1000 in PBS for nuclear staining. Cells were washed three times and wells were aspirated and mounted with coverslips using ProLong Diamond Antifade mountant (Thermo Fisher, Waltham, WA). Cells were imaged for GFP-positive puncta using a confocal Leica DMi8 microscope (Leica Microsystems) with oil immersion objective at 40X and 60X to report maximum intensity projections. Sample size was 4 per group.

For live-cell imaging, the reporter cells were seeded in the same way described above. Upon the LNP treatment, the cells were imaged for 16 hr with 30 min intervals for time-lapse live-cell imaging using Yokogawa CSU-X1 on Zeiss Axio Observer spinning disk microscope. The images were processed and reported as the maximum intensity projections in the manufacturer software (Zen, Zeiss), followed by additional processing in ImageJ for movie export.

### Puncta counting

Gal8-GFP puncta were quantified using ImageJ. Confocal images stacks were preprocessed using maximum intensity projections and Gaussian smoothing with radius of 1. Then, the puncta were counted using Find Maxima with prominence > 30. The puncta counts were normalized by the nuclei counts in the analysis, which were counted using standard segmentation procedures (automatic thresholding and watershed). For live-cell imaging, the puncta were counted in the analogous manner except the counts were normalized by the ratio (initial number of sample cells) / (initial number of control cells).

### Immunofluorescence (ICC)

At 3 hr and 24 hr timepoints after LNP-Chol and LNP-Sito transfections, cells were rinsed with PBS and fixed in 4% paraformaldehyde at room temperature for 10 minutes. Following fixation, cells were washed three times and blocking was performed for 2 hours at room temperature in PBS supplemented with 5% donkey serum and 0.2% Triton X-100 (Fisher Scientific, Waltham, MA). Cells were then rinsed with PBS and primary antibody dilutions for Rab7 (Cell Signaling, #9367) and EEA1 (Cell Signaling, #2411) were prepared at 1:100 in PBS supplemented with 1% bovine serum albumin (BSA) and 0.2% Triton X-100 (Fisher Scientific, Waltham, MA). Primary antibody was incubated overnight at 4°C with gentle rocking. The next day, the cells were washed with PBS three times and incubated with Alexa Fluor Plus 647-conjugated donkey anti-rabbit secondary antibody (Thermo Fisher, A32795) at 1:2000 in PBS supplemented with 1% BSA and 0.2% Triton X-100 for 2 hours at room temperature. Following incubation, cells were washed with PBS, nuclei stained with DAPI and coverslipped using ProLong Diamond Antifade mountant (Thermo Fisher, Waltham, MA). Cells were imaged using a Zeiss LSM 880 with Fast Airyscan employing a 63X oil immersed objective for maximum intensity projections reported. Sample size was 4 representative images per group. Image processing and export were done in the manufacturer’s software (Zen, Zeiss).

## RESULTS

### Characterization of LNPs consisting of various sterol analogs

In order to examine the effects of sterols in LNP trafficking in the endosomal pathway, we prepared a series of LNPs containing various sterol analogs. Cholesterol (Chol) was replaced with four different naturally occurring sterol analogs that have variations in the steroid backbone and the C24 side-chains. More specifically, β-sitosterol (Sito) and fucosterol (Fuco) have an ethyl group and an ethylidene group at C24-position, respectively. Campesterol (Camp) has a C24 methyl group and stigmastanol (Stig) is the reduced product of β-sitosterol that has a single bond at the Δ-5-position instead of a double bond (**Fig. 1B**). These phytosterols are known to alter cholesterol metabolism and the intracellular trafficking when incorporated in liposomes, indicative of their bioactivities.^29^ The micellar formulations of Sito and Camp were reported to alter cholesterol influx and secretion in cells.^30,31^ More importantly, the chemical diversity of the phytosterols can have effects on lipid packing when incorporated in LNPs owing to different stereochemistry, affecting the physicochemical properties of the nanoparticles. For example, Fuco has a more rigid side-chain at the C24 position as compared to Sito due to a double bond, and Stig has a highly flexible steroid ring because of reduction of the Δ−5 double bond. For these reasons, we hypothesized that substitution of sterol analogs in place of Chol in LNPs will change the nanoparticle characteristics, altering the endosomal trafficking and the incidence of endosomal escape. While the sterol components were varied, DLin-MC3-DMA, DSPC, and DMG-PEG_2K_ were used as the ionizable lipid, the structural lipid, and the PEG-lipid, respectively (**Fig. 1A**). LNPs containing Chol served the baseline formulation for comparisons throughout the study.

We first evaluated the physicochemical properties of a series of LNPs containing five sterol analogs (Chol, Sito, Fuco, Camp, and Stig) by dynamic light scattering (**Fig. 1B**). Particle sizes of LNPs containing Chol, Sito, Fuco, and Camp were similar (< 100 nm) whereas LNP-Stig showed a relatively larger size (> 110 nm) (**Fig. 1C**). Similarly, LNP-Chol, Sito, Fuco, and Camp were narrowly-distributed (PdI < 0.2) whereas the size distribution of LNP-Stig were relatively wide (PdI > 0.3) (**Fig. 1D**). In spite of size changes in LNP-Stig, all LNPs tested showed slightly negative zeta-potentials (**Fig. 1E**). The results of mRNA encapsulation assay showed that LNP-Stig had a relatively lower encapsulation efficiency (90%) while other LNPs (Chol, Sito, Fuco, and Camp) had more than 95% encapsulation efficiency (**Fig. 1F**).

Next, we investigated if the substitution leads to the morphological changes of LNPs using cryogenic transmission electron microscopy (cryo-TEM) (**Fig. S1**). Cryo-TEM analysis of LNP-Chol and -Camp showed spherical shapes whereas LNP-Sito and -Stig exhibited multi-faceted shapes (**Fig. S1**). LNP-Fuco displayed a distinctive morphology containing electron-dense pockets within a particle, which might be due to the hydrophobicity conflicts between LNP components (**Fig. S1**). These results concur with our previous reports that replacement of cholesterol with plant-based derivatives leads to the morphological changes.^32,33^

### Cellular entry of LNPs containing various sterol analogs

Having observed that changing physicochemical properties of the LNPs with varying sterols, we tested the cell uptake, cytotoxicity, and mRNA transfection of the LNPs in 293T/17 cells. To measure the cell entry of LNPs encapsulating mRNA, we used Cy5-labelled mRNA as a cargo. Cy5 intensity in the 293T/17 cells treated with various LNPs for 3 hr was measured using flow cytometry with a viability marker staining. We found that more than 98% single viable cells were Cy5-positive and average intensities (MFI) of Cy5 within the cells were not altered by various sterol substitutions in LNP formulations, indicating that endocytosis of various LNPs were similar regardless of sterol analogs used (**Fig. S2**).

### Gal8 visualization indicating LNP-mediated endosomal disruption

Next, we evaluated Gal8 recruitment upon LNP treatment, suggestive of the incidence of endosomal escape events. This redistribution of Gal8 in response to the endosome damage sensing is visualized as GFP puncta in the Gal8-GFP reporter cells. Therefore, we counted the GFP puncta for cells treated with our panel of LNPs to attempt to visualize differences of the elusive endosomal escape events of these nanoparticles.

We looked at Gal8 recruitment in cells at 3 and 24 hr treatment with LNPs. Representative images of each LNP treatment are shown in **Fig. 2A, Fig. S3 and S4**. All LNPs led to some level of recruitment of Gal8-GFP in the reporter cells. Increasing mRNA dose produced more counts of Gal8 puncta and the overall puncta counts were higher, and more readily detectable in the 24 hr treatment compared to 3 hr treatment (**Fig. 2B, C**). After 3 hr incubation, LNP-Fuco produced significantly more Gal8 puncta when compared to LNP-Chol in all mRNA doses tested (**Fig. 2B**). LNP-Camp and -Stig also produced high degrees of Gal8 recruitment whereas LNP-Chol and -Sito did not change the puncta counts in 3 hr treatment (**Fig. 2B**). After 24 hr treatment, LNP-Fuco, -Camp, and -Stig produced significantly higher degrees of easy to detect Gal8 recruitment in all mRNA doses tested (**Fig. 2C**). LNP-Sito also increased the Gal8 recruitment in the doses of 100 ng/well and 200 ng/well whereas LNP-Chol induced mild Gal8 recruitment only at 200 ng/well (**Fig. 2C**). Despite the weaker recruitment in 3 hr results, both timepoints demonstrated the potential of this Gal8-GFP system to cause a large amount of easily detectable puncta for the elucidation of rare endosomal escape events.

**Fig. 2.**
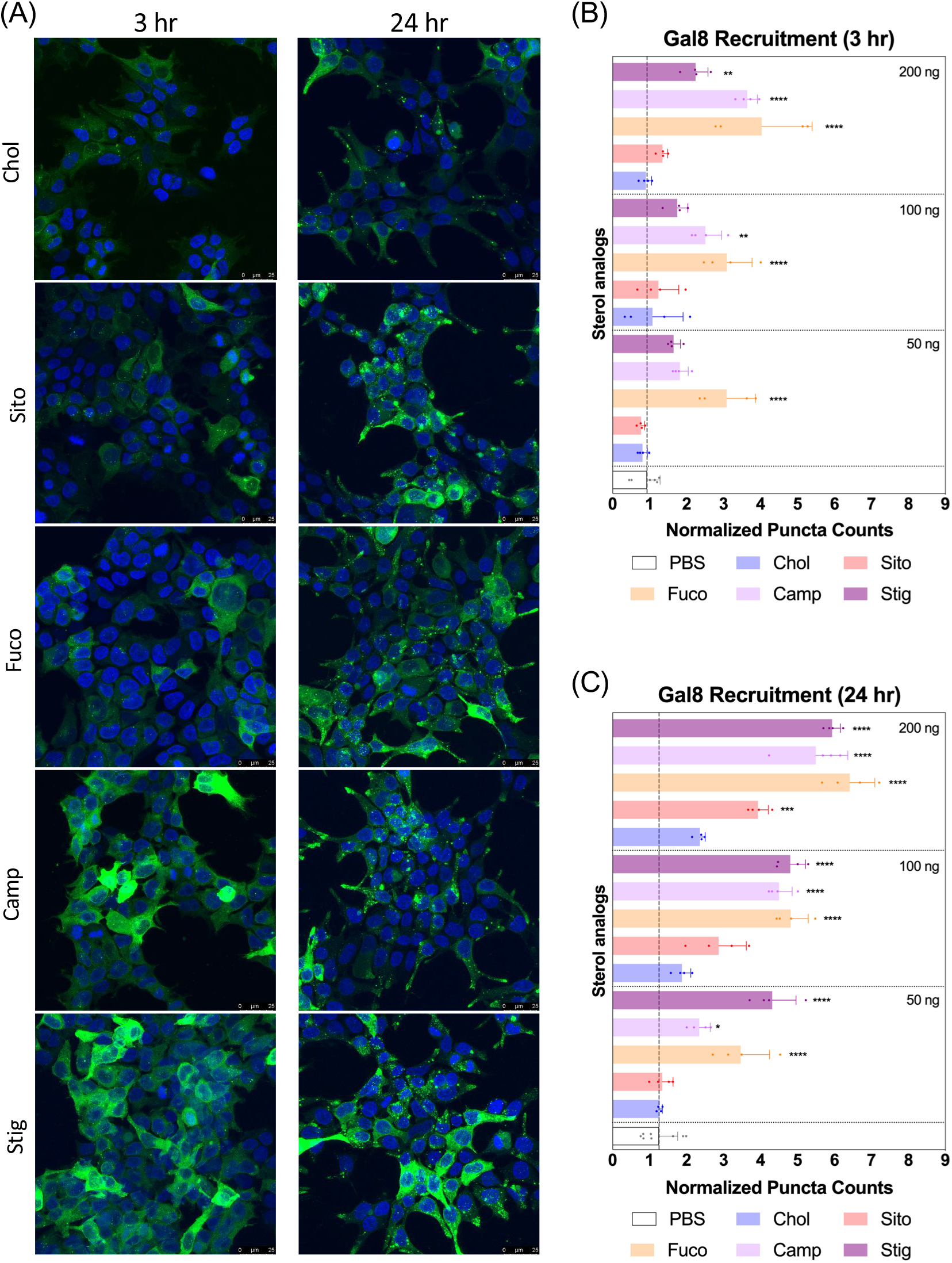
**(A)** Representative images of the Gal8-GFP reporter cells after treatment of the LNPs containing various sterol analogs for 3 and 24 hr. Presented in maximum intensity projection. Gal8-GFP (green) and nucleus (blue). Scale bars show 25 μm. (B, C) Normalized puncta counts indicating Gal8 recruitment in the reporter cells after treating with the LNPs for **(B)** 3 hr and **(C)** 24 hr. mRNA doses given were displayed on the right corners of each segment: 50, 100, and 200 ng per well. Statistical analyses were performed against LNP-Chol at each dose using Tukey’s multiple comparisons test (n = 4); *p<0.05; **p<0.01; ***p<0.001; ****p<0.0001

### mRNA transfection of LNPs containing sterol analogs

Next, we evaluated *Fluc* mRNA transfection by various LNPs in 293T/17 cells. We assayed the cell viability and the luciferase expression at two different timepoints: 3 and 24 hr for LNP treatment (**Fig. 3**). The cell viability was not compromised by any treatment of various LNPs at any of the doses tested (**Fig. 3A, B**). Luciferase assay results showed that all LNPs produced similar levels of luminescence at the doses of 50 and 100 ng/well. At 200 ng/well, the highest dose employed, LNP-Sito, -Fuco, -Camp resulted in greater luciferase expressions than LNP-Chol while LNP-Stig produced the lowest expression in all three doses tested (**Fig. 3C**). Influence of sterol substitutions on mRNA transfection became more pronounced in the 24 hr treatment. Similarly to the results from 3 hr treatment, LNP-Sito, -Fuco, and -Camp outperformed the mRNA transfection when compared with LNP-Chol (**Fig. 3D**) in the 24hr data collected. In particular, LNP-Camp significantly enhanced mRNA transfection in all doses while LNP-Sito and -Fuco did so in the 200 ng/well condition as compared to LNP-Chol. LNP-Stig showed poor mRNA transfection ability at all mRNA doses tested. Taken together, varying luciferase expression is likely due to the different rates of endosomal escape given that the cellular uptake of various LNPs was similar. The lack of correlation between mRNA expression and Gal8 puncta counts of LNP-Stig will need to be further explored as it is an unexpected, not yet elucidated result.

**Fig. 3.**
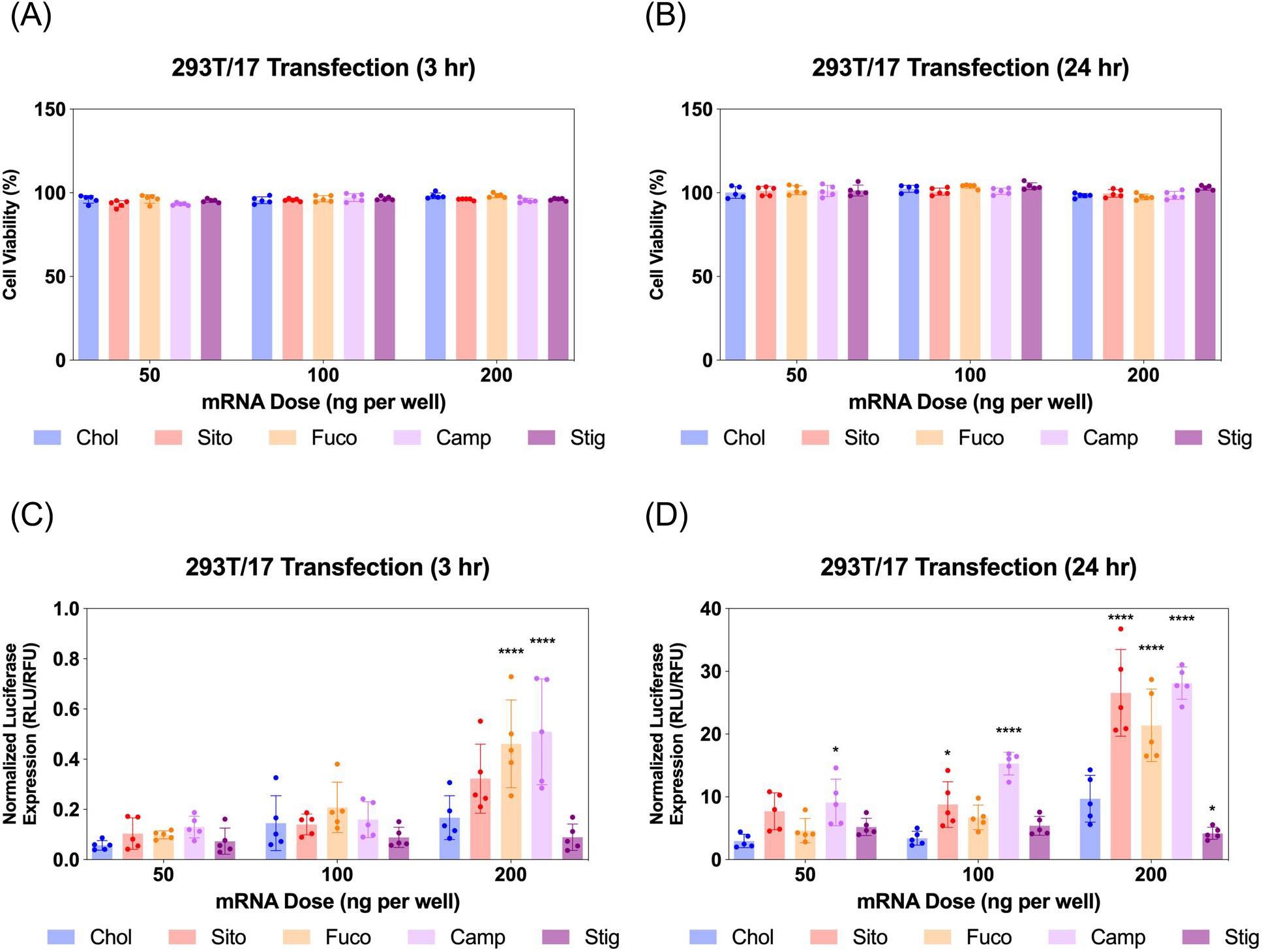
mRNA transfection assay in 293T/17 cells treated with LNPs containing various sterol analogs: Chol (blue), Sito (red), Fuco (orange), Camp (lavender), Stig (purple) **(A, B)** Cell viability of 293T/17 cells treated with the LNPs containing various sterol analogs for **(A)** 3 hr and **(B)** 24 hr. **(C, D)** Luciferase expression in the transfected 293T/17 cells using the LNPs containing various sterol analogs for **(C)** 3 hr and **(D)** 24 hr.

### Live imaging of LNP-driven endosome damage inside Gal8-GFP reporter cells

Differential Gal8 recruitment with sterol substitutions in LNPs led us to focus on the LNP-Chol and our well characterized lead formulation, LNP-Sito. In conjunction with the moderate increase by LNP-Sito in Gal8 puncta counts, our previous findings demonstrating that Sito substitution (referred as eLNP in the previous reports) alters the morphology,^33^ diffusivity in endosomes, and endosomal trafficking^32,34^ of LNPs and enhances mRNA transfection *in vitr*o^32,33^ and *in vivo*^35^ reinforced the decision to pick these two LNPs to test further. To exclude any possible artifact coming from fixation, we performed time-lapse live-cell imaging using spinning disk confocal microscopy. Representative images and time-lapse movies of control, LNP-Chol and LNP-Sito are provided in **Fig. 4A** and supplemental videos (**Supporting Movie 1-3**). Both LNP-Chol and -Sito produced Gal8 recruitment; while control group did not show any recruitment (**Fig. 4A**). Notably, LNP-Sito strongly induced the Gal8 recruitment, resulting in ten-fold higher number of puncta than LNP-Chol throughout the imaging (**Fig. 4A, B**). Within 30 min after the LNP treatment, the Gal8 reporter cells rapidly reacted to the LNPs and produced GFP puncta (**Fig. 4B**). The normalized punctae count of LNP-Sito outnumbered that of LNP-Chol during 16 hr live-cell imaging (**Fig. 4B**). Additionally, Gal8 recruitment induced by LNP-Sito showed an increasing trend in the course of time whereas that of LNP-Chol remained relatively constant, congruent with previous baseline-like results (**Fig. 4B**). This striking difference in the GFP punctum population suggests the highly efficient endosomal escape of LNP-Sito, which corresponds to the mRNA expression results (**Fig. 2C, E**).

**Fig. 4.**
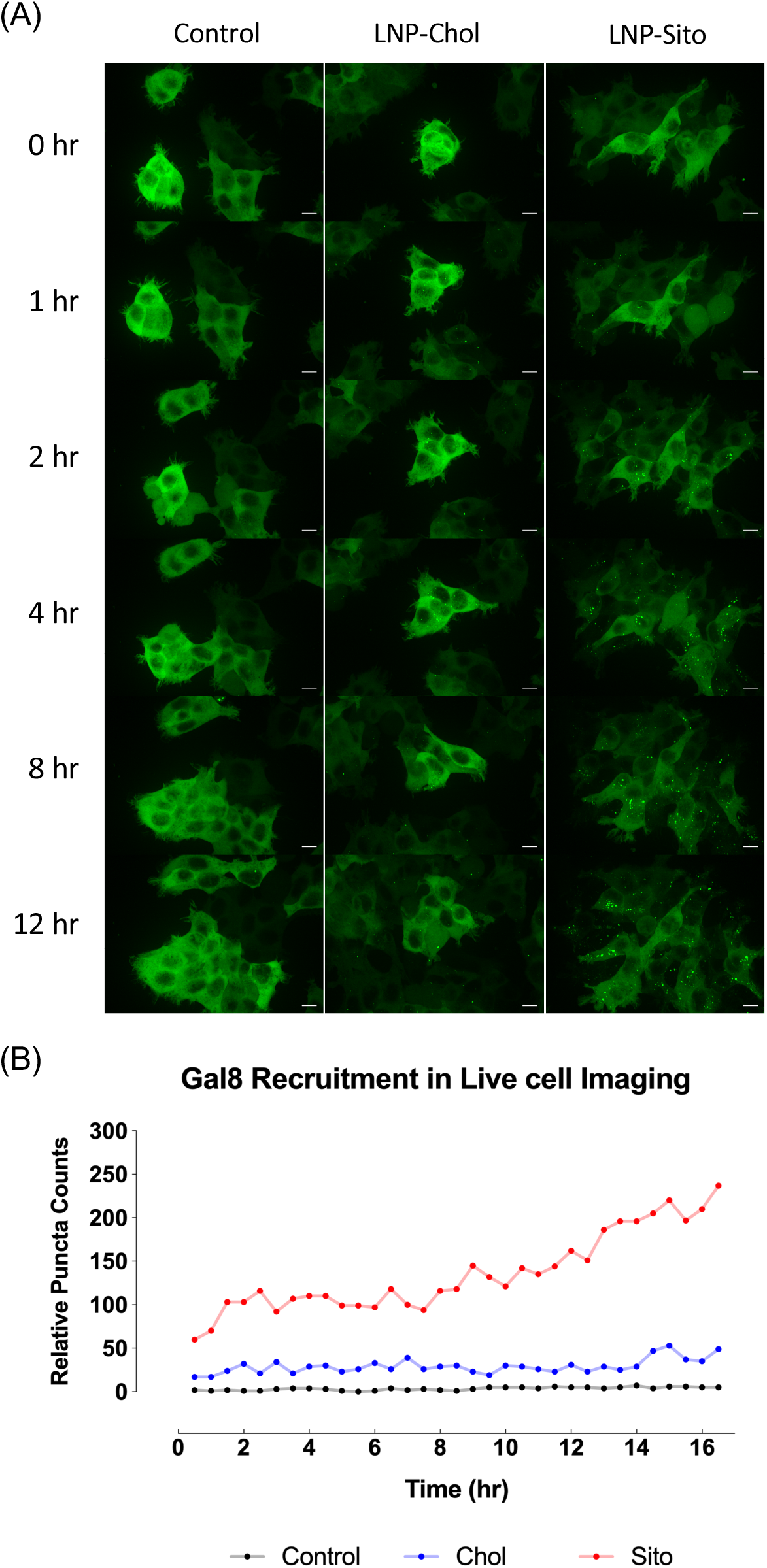
**(A)** Representative snapshots from live-cell imaging of the Gal8-GFP reporter cells after treatment of serum-free media (Control), LNP-Chol and LNP-Sito (at 100 ng mRNA per well). Presented in maximum intensity projection. Scale bars show 10 μm. **(B)** Relative puncta counts in the reporter cells after treatment of serum-free media (Control, black), LNP-Chol (blue) and LNP-Sito (red). LNPs were treated at a dose of 100 ng mRNA per well. Time-lapsed snapshots were taken every 30 min for 16 hr. Z-stack snapshots were processed for maximum intensity projection for Gal8 puncta counting.

### Subcellular colocalization analysis of Gal8 recruitment

To interrogate which stage of the endosomal pathway is associated with Gal8 recruitment, we conducted immunofluorescence with the endosome markers: EEA1 (early endosome) and Rab7 (late endosome) (**Fig. 5**). We also quantified the colocalization of Gal8 puncta and the endosome markers by calculating the Pearson’s correlation coefficient (PCC) and the Mander’s correlation coefficient (MCC) (**Fig. S5**). In overlaid image analysis, GFP puncta representing the Gal8 recruitment were not well colocalized with the EEA1 staining, which was confirmed in the correlation coefficients (**Fig. 5A, Fig. S5A**). Both PCC and MCC analyses indicate that Gal8 puncta had no positive-correlation with the EEA1 staining (e.g., approximately 0 for LNP-Chol and 0.05 for LNP-Sito in MCC), suggesting that Gal8 recruitment, with associated endosome disruption, is not happening in the early endosomes (**Fig. S5A**). On the other hand, many of the Gal8 puncta overlapped with the Rab7 staining as evident both from overlaid images (**Fig. 5B**) and metric analysis (e.g., approximately 0.2 for LNP-Chol and 0.6 for LNP-Sito in MCC) (**Fig. S5B**). Particularly, LNP-Sito produced the higher PCC and MCC than LNP-Chol, which is perhaps due to the stronger recruitment of Gal8 by LNP-Sito (**Fig. S5**). Overall, better colocalization of Gal8 puncta with Rab7 than EEA1 for both LNP-Chol and -Sito suggests that LNPs disrupted the late endosomes to enter the cytosol.

**Fig. 5.**
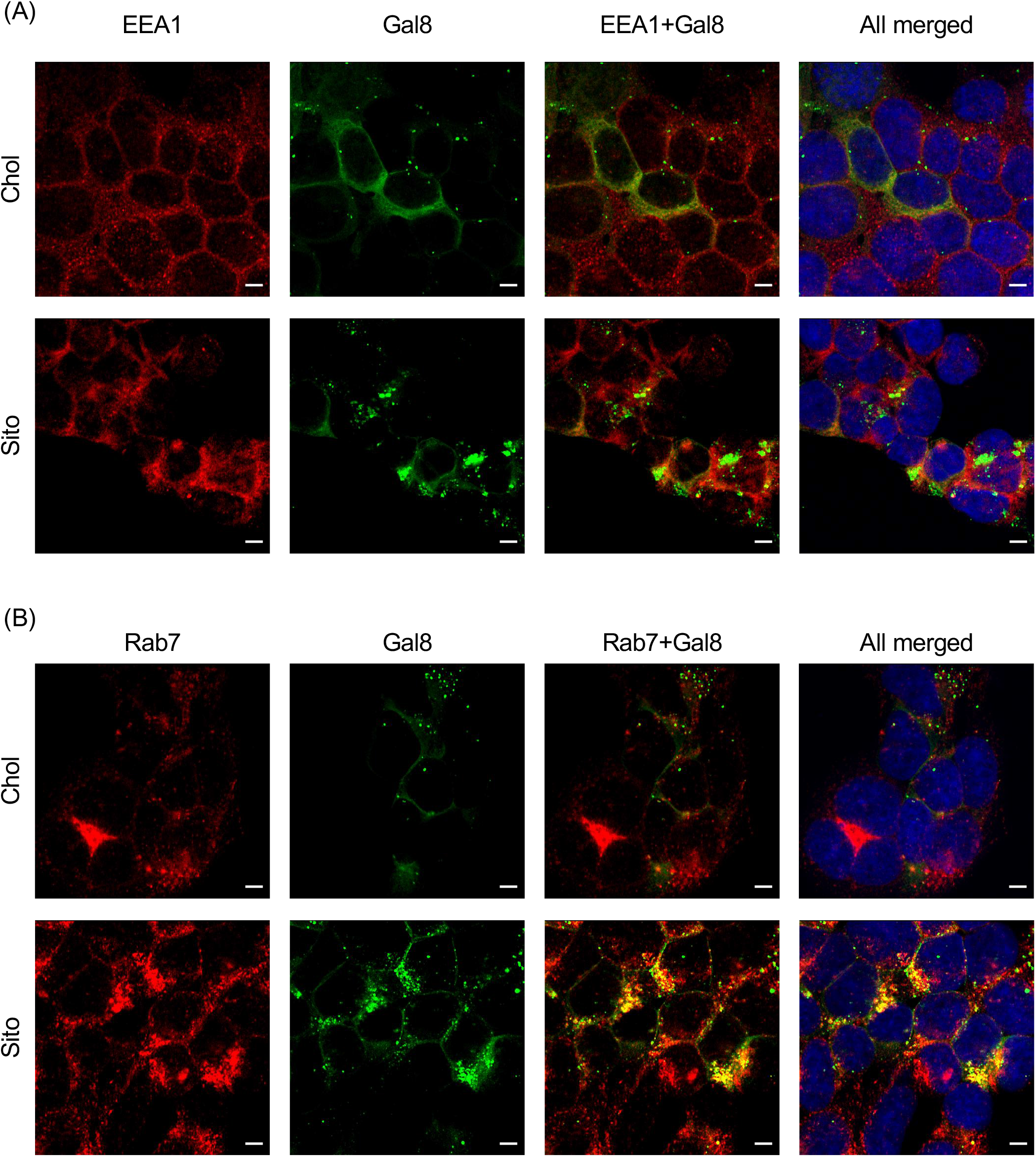
**(A, B)** Representative images of the Gal8-GFP reporter cells stained with endosomal markers: **(A)** EEA1 (early endosome) and **(B)** Rab7 (late endosome). Red: endosomal markers, green: Gal8-GFP, blue: nucleus. Scale bars show 5 μm.

## DISCUSSION

LNPs are sophisticated nanocarriers made up of multiple elements: an ionizable lipid, helper lipids (namely cholesterol and phospholipid), and a PEG-lipid (**Fig. 1A**). The ionizable lipid has gained a lot of attention because of its primary roles in RNA complexation, cellular uptake, and release of the genetic cargo, resulting in a lot of studies to discover novel ionizable lipids.^36–40^ In contrast, the effects on bioactivity of cholesterol is less explored in spite of being the second largest content in the clinically approved LNP formulation.^36,41^ Cholesterol in the LNPs modulates fluidity and permeability of lipid membrane by enhancing lipid packing.^32^ It is conceivable that chemical modifications to cholesterol influence these effects. For instance esterified cholesterol variants enhanced LNP-mediated gene delivery whereas oxidized counterparts did not show any change.^42^ Our group also recently demonstrated that substitution of cholesterol with phytosterols in the LNPs influences nanocarrier morphology, internal structures, and transfection efficiency, forecasting changing numbers of disrupted endosomes inside the cell.^32,33^ Despite encouraging findings, the molecular web connecting LNP morphology to endosomal escape events and subsequent localization of nucleic acids in the cytosol remains murky, especially in the context of direct detection of endosome destabilization events.^12,43^

We have employed Gal8-GFP reporter cell system to visualize the endosome destabilization induced by treating with LNPs and found that sterol substitution in LNPs changes not only Gal8 recruitment but also translation of delivered mRNA (**Fig. 2A, 3D**). These findings suggest that sterol substitution in LNPs causes endosomal escape at varying levels, affecting the mRNA transfection efficiency. Accompanying the clear difference in Gal8 punctum populations between the LNPs containing sterol analogs substituted, the level of Gal8 recruitment correlated well with the mRNA transfection efficiency of most except for one of our LNPs tested. LNP-Stig, which strongly recruited Gal8 puncta, exhibited the poorest mRNA transfection in the parental 293T/17 cells. This unexpected discrepancy between luciferase expression and the normalized puncta counts could be due to suboptimal choices in treatment time or mRNA doses specific to each formulation used, or possible artifacts stemming from fixation. Otherwise, it could be due to the cell type-specificity or batch-to-batch variation of Stig since our previous reports with LNP-Stig showed effective mRNA transfection in HeLa cells. Further investigation of discrepancies observed may provide valuable clues regarding the LNP-incorporated cholesterol sensing in the endosomal pathway, the cytosol and the complex other machineries involved along this escape and subsequent translation of cargo.

Live-cell imaging of LNP-Chol and -Sito evinced a visually robust temporal variation of endosome disruption depending on sterol type incorporated in LNPs. LNP-Sito, (also known as eLNP) showed highly notable levels of Gal8 puncta during 16 hr live-cell imaging. Given that extended retention of LNP-Sito than LNP-Chol inside endocytic vesicles,^32^ one could speculate this could lead LNP-Sito to potentially effect more efficient endosomal escape, generating the greater number of Gal8 puncta observed. The high retention of LNP-Sito could have boosted the endosome escape by accumulating in the endosomes at the later timeframes (after 13 hr, **Fig. 4B**). This reasoning is reinforced by the evidence showing intracellular delivery of nucleic acids is enhanced by extended retention of LNPs at the late endosomes and lysosomes in Niemann-Pick disease, type C1 (NPC1) knockout cells.^12^ Therefore, if NPC1 and 2 handle Sito differently from Chol, LNP-Sito could get extra time in the late endosomes, leading to more endosomal destabilization and escape.

Colocalization with endosome markers revealed the spatial distribution of Gal8 recruitment in the context of the endosomal pathway. Our results show that endosomal escape of LNPs is not colocalized well with the early endosomes (**Fig. 5A, S5A**), but preferentially occurs in late endosome (**Fig. 5B, S5B**). This is corroborated in a previous report by Wittrup et al. showing that Gal8 recruitment mediated by siRNA-loaded LNPs does not occur at EEA1-positive endosomes, but at Rab5- or Rab7-positive endosomes.^17^ In agreement with how reported Gal8 sensing to damaged endosomes leads to the development of autophagosomes, which fuse with lysosomes for degradation or exocytosis,^11^ colocalization of Gal8 puncta and lysotracker in the study of Kilchrist et al. indicates a portion of damaged endosomes being transported to lysosomes.^26^ Another study by our group also highlights the importance of late endosome development for successful mRNA delivery, as supported by the observation that LNP-mediated mRNA transfection was suppressed in Rab7-knockout cells.^34^ Gal8 is also involved in signaling of mTOR (mechanistic target of rapamycin) which is recruited to late endosomes and lysosomes for initiating cell proliferation events, such as protein synthesis and autophagy, when cells are supplied with nutrients.^44^ Our group previously showed that mTORC1 (mTOR complex 1) inhibitors decrease the translation of delivered mRNA due to diminished protein synthesis, while constitutive activation of mTORC1 by genetic deletion of tuberous sclerosis complex 2 (TSC2), which is a negative regulator of mTORC1, increases the translation of delivered mRNA.^34^ Conversely, Gal8 recruitment to the damaged endosomes inactivates mTORC1, inhibiting the cell proliferation events.^44^ These conflicts may be explained by perhaps quantitative easing induced by a large number of mRNA molecules escaping into the cytosol in spite of reduced mTORC1 signaling due to the endosome disruption. Although speculative, these hypotheses underline the highly complex molecular dynamics that are at play before, during and immediately after endosomal escape of nanoparticles. Taken together, our results support that the primary site of LNP-mediated endosomal escape is ranged from late endosomes to lysosomes, which can be further enhanced by sterol substitution.

## CONCLUSIONS

In sum, we confirmed that the cholesterol in LNPs has pronounced effects on morphology, internal structures and mRNA transfection efficiency. The massive endosomal escape produced with C24 alkyl derivatives of cholesterol within LNPs can lead to identification of vesicles that were previously hard to pinpoint due to rarity of the escape events. Late endosomal biogenesis and trafficking emerge, once again, as the site of interest for elucidation of LNP endosomal escape. We observed stronger colocalization of LNP escape events with the Rab7 late endosomal marker than with the EEA1 early marker. Future studies should concentrate around other endosomal stage markers in conjunction with knockout cell lines of other involved players to dissect the spatiotemporal intricacies of endosomal trafficking and successful escape into the cytoplasm. Further characterization of the molecular interactions of LNP components with the endolysosomal membrane will allow us to crack some of the puzzling findings reported for LNPs containing C24 sterol derivatives. Shedding light on the structure-activity relationships of these sterol components with their interacting moieties will allow for the rational engineering of the next generation LNPs. Other signaling moieties such as other galectins could be of interest especially considering the complex regulatory roles they effect in the endolysosomal system as a whole. Additionally, expanding our understandings to *in vivo* areas is of greatest importance to empower nanomaterials for gene therapy. In conclusion, we advocate for the concerted probing of these known unknowns so that we can contribute to the breakthrough of this biological labyrinth.

## Supporting information

Electronic supplementary information

## ACKNOWLEDGEMENTS

This project was supported through funding from the National Heart Lung and Blood Institute (N.H.L.B.I) 5R01HL146736 (G.S), National Eye Institute 1R21EY031066 (G.S), and Cystic Fibrosis Foundation SAHAY19XX0 (G.S). A portion of this research was supported by NIH grant U24GM129547 and performed at the Pacific Northwest Cryo-EM Center (PNCC) at OHSU and accessed through EMSL (grid.436923.9), a DOE Office of Science User Facility sponsored by the Office of Biological and Environmental Research. We thank Dr. Stefanie Kaech Petrie, Dr. Brian Jenkins, and Dr. Crystal Chaw in the Advanced Light Microscopy Core in OHSU for the help for live-cell imaging. We thank the flow cytometry core in OHSU for the help for cell sorting.

## FOOTNOTES

Electronic supplementary information (ESI) available: Cryo-TEM analysis and cellular uptake of LNPs, representative images of Gal8-GFP reporter cells treated with LNPs at various doses, colocalization analyses of Gal8 puncta with endosomal markers, time-lapse movies recording live-cell imaging of Gal8-GFP reporter cells treated with serum-free media, LNP-Chol, or LNP-Sito.

