## Supplementary material for "Illuminating endosomal escape of polymorphic lipid nanoparticles that boost mRNA delivery": Electronic supplementary information

**Cryo-TEM image acquisition and processing**

Cryo-TEM acquisition was performed at 300 kV using FEI Titan Krios (Thermo Fisher, Hillsboro, OR) equipped with Falcon III and K3 cameras with DED. 3-5 ul of the sample was dispensed on a plasma cleaned grid (Quantifoil, R1.2/1.3 300 or 400 Cu mesh) in the FEI Vitrobot chamber at 95% relative humidity and allowed to rest for 30 seconds. Then, the grid was blotted for 3 seconds with filter paper and plunged into liquid ethane cooled by liquid nitrogen. The frozen grids were then checked for visible defects and assembled into cassettes, and the acquired images were analyzed using ImageJ. The images were processed using low-pass filter.

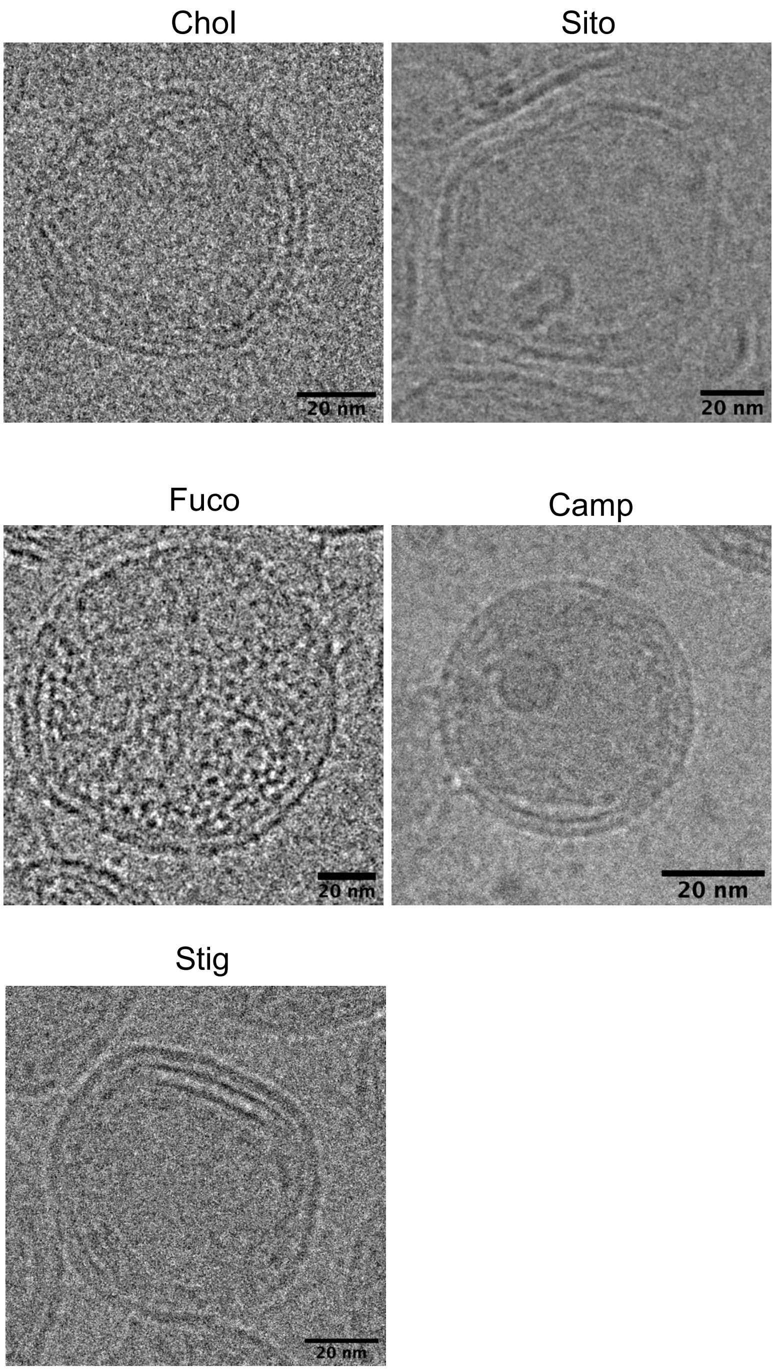


**Fig. S1.** cryo-TEM micrographs of the LNPs containing various sterol analogs. Scale bars show 20 nm.

**Evaluation of LNP cellular uptake using flow cytometry**

293T/17 cells were seeded at 100,000 cell per well in 6-well plates, followed by overnight incubation for cell adhesion. The cells were treated with the LNPs encapsulating Cy5-labelled EGFP mRNA at 500 ng mRNA per well for 3 hr. Then, the cells were processed to single cell suspensions using TrypLE Express (Thermo Fisher), followed by staining with Fixable Viability Dye eFluor$^{TM}$ 780 and fixing with 1% paraformaldehyde. The samples were diluted with sterile PBS and analyzed in BD Fortessa with 670/30 and 780/60 filters. The data were processed in FlowJo software v10.7 (FlowJo LLC, Ashland, OR) to gate populations and to calculate mean fluorescence intensity of Cy5.

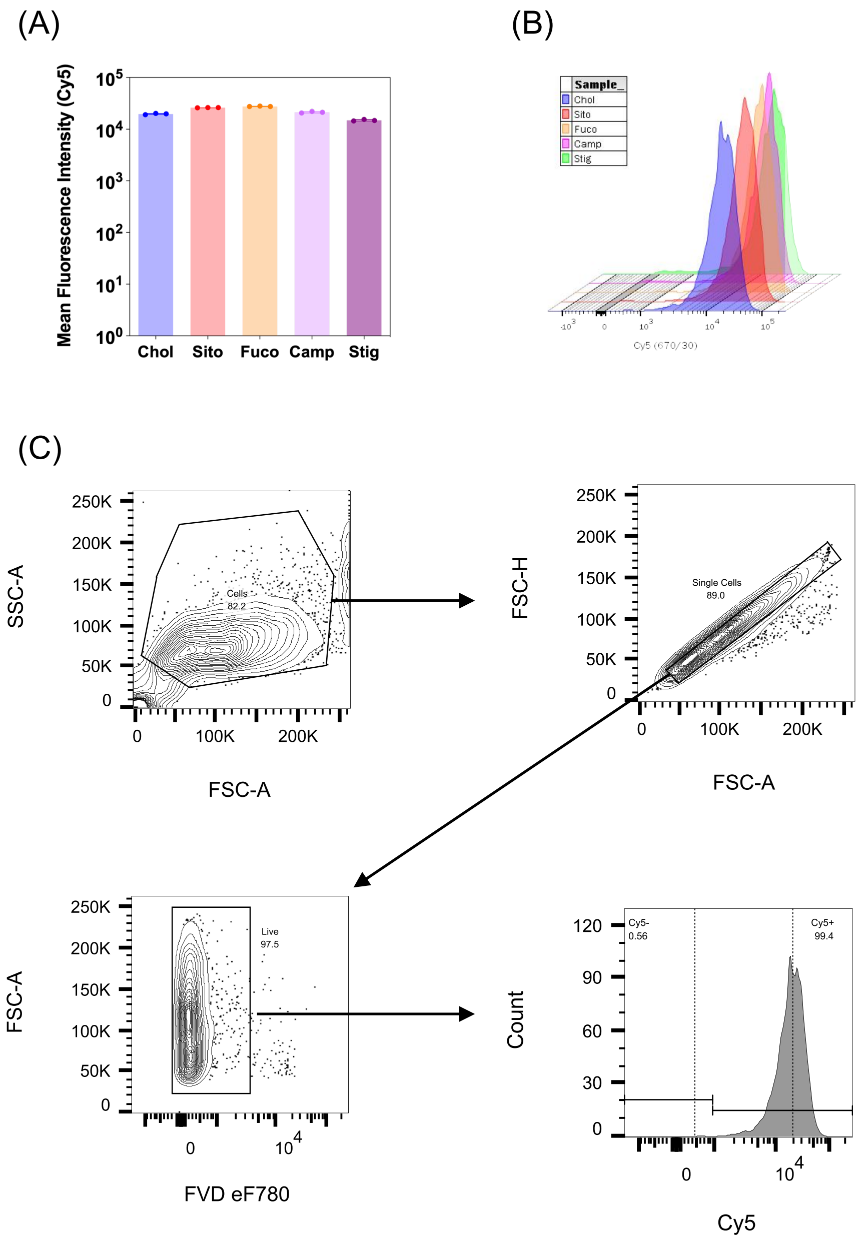


**Fig. S2. (A)** Cellular uptake of LNPs containing various sterol analogs (n = 3). **(B)** Multigraph overlay of Cy5 histograms of the samples treated with LNPs containing various sterol analogs: Chol (blue), Sito (red), Fuco (orange), Camp (pink), and Stig (green). **(C)** Gating strategy for Cy5-positive cell populations.

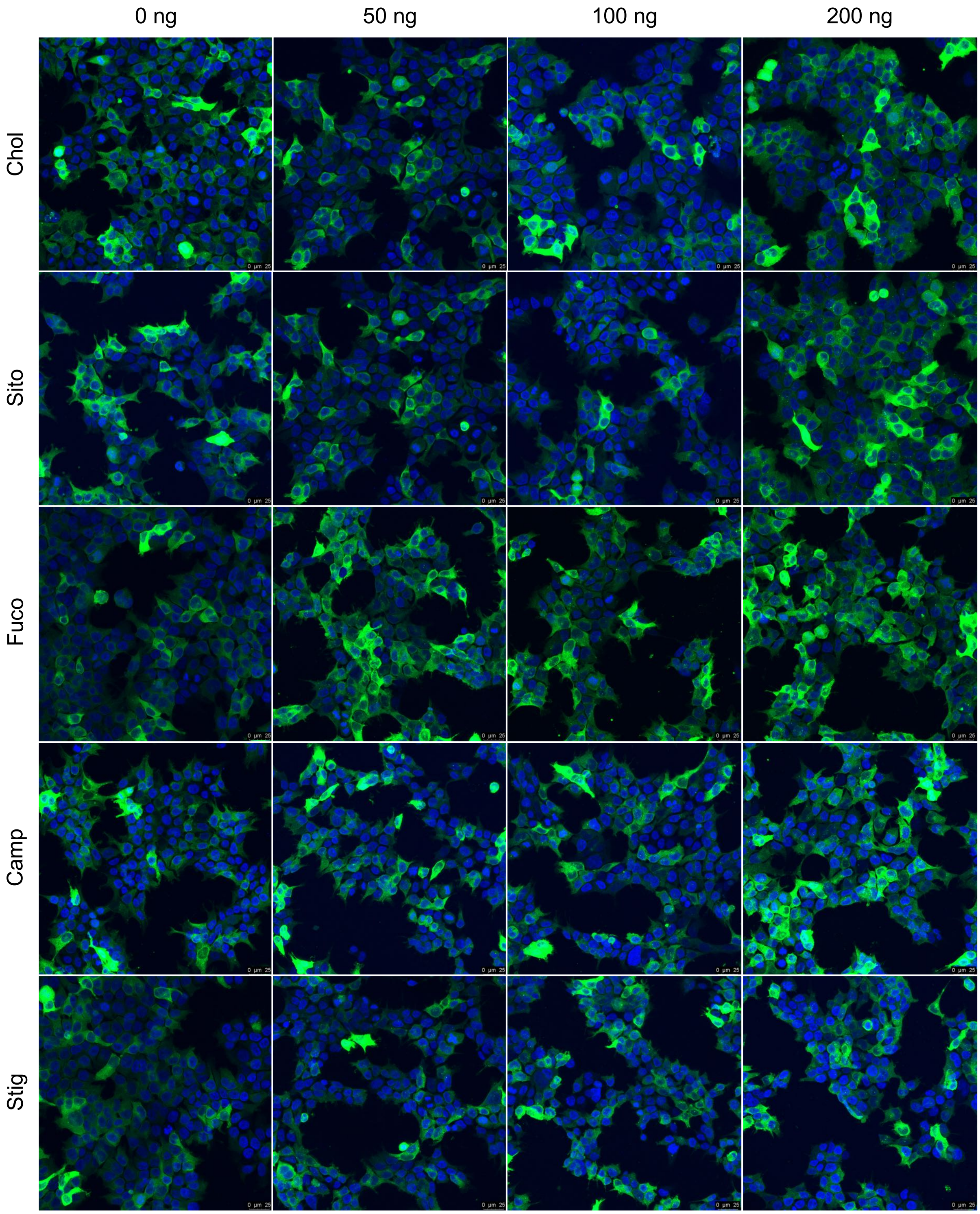


**Fig. S3.** Representative images of the Gal8-GFP reporter cells after treatment of the LNPs containing various sterol analogs at various mRNA doses for 3 hr. Presented in maximum intensity projection. Gal8-GFP (green) and nucleus (blue). Scale bars show 25 µm.

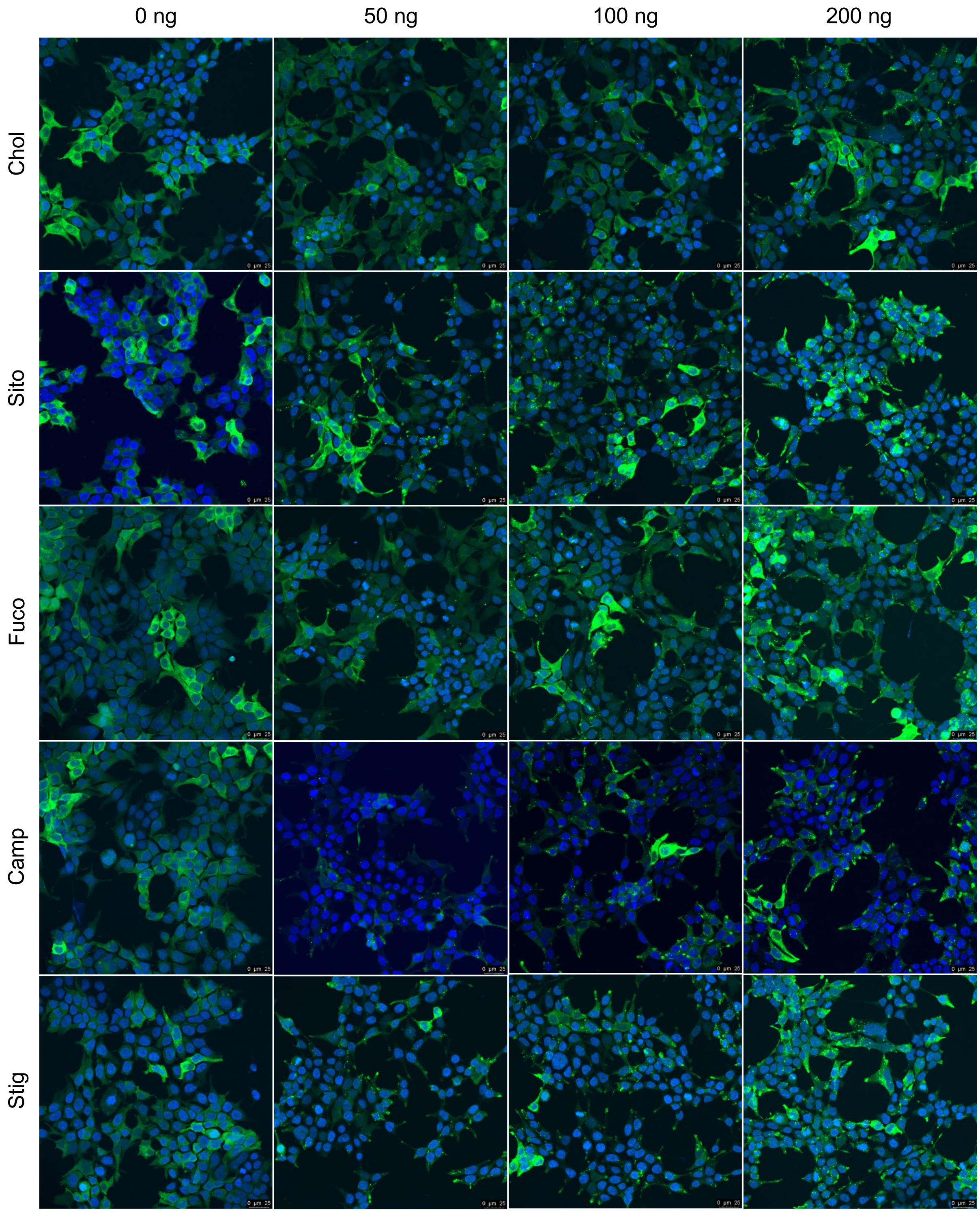


**Fig. S4.** Representative images of the Gal8-GFP reporter cells after treatment of the LNPs containing various sterol analogs at various mRNA doses for 24 hr. Presented in maximum intensity projection. Gal8-GFP (green) and nucleus (blue). Scale bars show 25 µm.

### Colocalization analysis

Colocalization analysis was performed on maximum intensity Z-projections of confocal image stacks using Coloc2 plugin of ImageJ. In order to disregard the cytosolic GFP signal, we only selected the areas with puncta in GFP-Gal8 channel by applying a mask. To create the mask, we selected the maxima with prominence > 3000 and created the mask with maxima points within tolerance. Pearson's (before automatic bisection thresholding) and Mander's correlation coefficients (after automatic thresholding) were selected for the discussion.

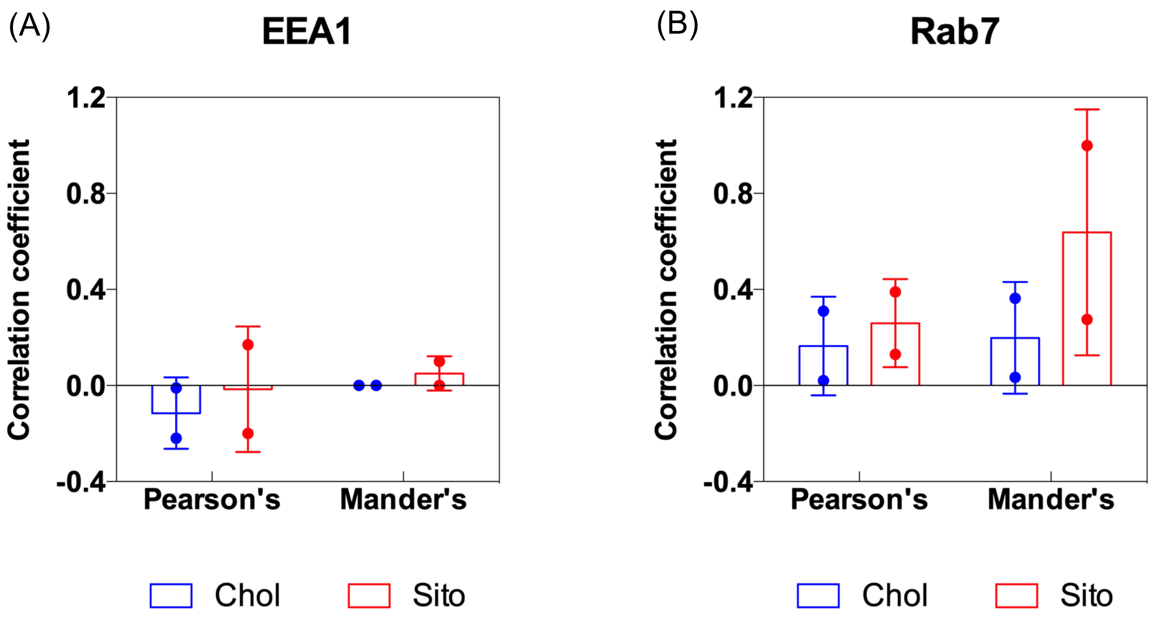


**Fig. S5.** Measuring colocalization of Gal8 puncta with endosomal markers, **(A)** EEA1 for early endosomes and **(B)** Rab7 for late endosomes. The Gal8-GFP reporter cells were treated with LNP-Chol (blue) or LNP-Sito (red) for 24 hr, followed by immunocytochemistry staining with EEA1 and Rab7 antibodies. The resulting images were processed to maximum intensity projection, followed by Pearson's and Mander's correlation analyses to calculate correlation coefficients in ImageJ (n = 2).
